# Interleukin-6 induces nascent protein synthesis in human DRG nociceptors via MNK-eIF4E signaling

**DOI:** 10.1101/2024.04.04.588080

**Authors:** Molly E. Mitchell, Gema Torrijos, Lauren F. Cook, Juliet M. Mwirigi, Lucy He, Stephanie Shiers, Theodore J. Price

**Affiliations:** Center for Advanced Pain Studies, School of Behavioral and Brain Sciences, University of Texas at Dallas, Richardson, Texas 75080

**Keywords:** nociceptors, human DRG, cytokine signaling, IL-6, MNK, eIF4E, azidohomoalanine

## Abstract

Plasticity of dorsal root ganglion (DRG) nociceptors in the peripheral nervous system requires new protein synthesis. This plasticity is believed to be responsible for the physiological changes seen in DRG nociceptors in animal models of chronic pain. Experiments in human DRG (hDRG) neurons also support this hypothesis, but a direct observation of nascent protein synthesis in response to a pain promoting substance, like interleukin-6 (IL-6), has not been measured in these neurons. To fill this gap in knowledge, we used acutely prepared human DRG explants from organ donors. These explants provide a physiologically relevant microenvironment, closer to *in vivo* conditions, allowing for the examination of functional alterations in DRG neurons reflective of human neuropathophysiology. Using this newly developed assay, we demonstrate upregulation of the target of the MNK1/2 kinases, phosphorylated eIF4E (p-eIF4E), and nascently synthesized proteins in a substantial subset of hDRG neurons following exposure to IL-6. To pinpoint the specific molecular mechanisms driving this IL-6- driven increase in nascent proteins, we used the specific MNK1/2 inhibitor eFT508. Treatment with eFT508 resulted in the inhibition of IL-6-induced increases in p-eIF4E and nascent proteins. Additionally, using TRPV1 as a marker for nociceptors, we found that these effects occurred in a large number of human nociceptors. Our findings provide clear evidence that IL-6 drives nascent protein synthesis in human TRPV1+ nociceptors via MNK1/2-eIF4E signaling. The work links animal findings to human nociception, creates a framework for additional hDRG signaling experiments, and substantiates the continued development of MNK inhibitors for pain.

## INTRODUCTION

Chronic pain affects the lives of billions across the world with patients suffering from debilitating spontaneous pain, hypersensitivity, and many comorbidities, yet current analgesic treatments remain largely ineffective (Finnerup et al., 2015; Dahlhamer et al., 2018; Mills et al., 2019; Price and Ray, 2019). The development and maintenance of chronic pain physiology is believed to be rooted in neuroplasticity with these changes occurring along the neuraxis. Important sites of neuroplasticity driving chronic pain are the DRG, the spinal dorsal horn, and multiple brain regions (Price and Inyang, 2015). In nociceptors, the sensory neurons responsible for detecting noxious stimuli, a key plasticity mechanism is an activity-dependent translation of new proteins from existing pools of mRNAs expressed by these neurons (Jimenez-Diaz et al., 2008; Geranton et al., 2009; Melemedjian et al., 2010; Melemedjian et al., 2014; Moy et al., 2017; Megat et al., 2019; Barragan-Iglesias et al., 2020). This process plays a crucial role in shaping the sensitivity and responsiveness of nociceptors in particular in response to growth factors like nerve growth factor (NGF) and cytokines like IL-6 (Melemedjian et al., 2010; Moy et al., 2017). This activity-dependent translation, which can be directly measured using nascent protein synthesis assays (Dieterich et al., 2006; Dieterich et al., 2007), can increase the excitability of nociceptors creating peripheral nociceptive signals that are ultimately perceived as pain (Moy et al., 2017; Megat et al., 2019; Mihail et al., 2019; Li et al., 2023).

Translation of mRNA is comprised of three steps: initiation, elongation, and termination. Initiation is considered the rate-limiting step for translation and is the step that is regulated most tightly in neurons when activity-dependent translation is stimulated (Kelleher et al., 2004; Costa-Mattioli et al., 2009; Yousuf et al., 2021). Extracellular signals such as NGF and IL-6 in DRG neurons, and neurotransmitters in CNS neurons can activate two major pathways to control activity-dependent translation. The major pathways are the mechanistic target of rapamycin (mTOR) pathways and the MAP kinase-interacting kinase (MNK1/2)/eukaryotic initiation factor (eIF) 4E, p-eIF4E pathway. (Costa-Mattioli et al., 2009; Khoutorsky and Price, 2018; Yousuf et al., 2021). Studies in rodent DRG neurons suggest that while both pathways play a role in regulating the excitability of nociceptors, MNK1/2-eIF4E signaling is important for controlling the excitability of these neurons (Moy et al., 2018a; Megat et al., 2019; Barragan-Iglesias et al., 2020; Jeevakumar et al., 2020; Shiers et al., 2020b), and is likely a more tractable pharmacological target for pain treatment for many reasons that have been described extensively elsewhere (Melemedjian et al., 2013; Khoutorsky et al., 2015; Megat and Price, 2018; Yousuf et al., 2021). Importantly, MNK1/2 is expressed in human nociceptors (Shiers et al., 2023) and its inhibition reduces the excitability of spontaneously active human DRG neurons recovered from patients with neuropathic pain undergoing thoracic vertebrectomy surgery (Li et al., 2023).

This study aimed to fill a gap in knowledge between a decade of studies in rodents linking IL-6*-* induced activity-dependent translation via MNK1/2-eIF4E signaling to enhanced nociception (Melemedjian et al., 2010; Melemedjian et al., 2014; Moy et al., 2017; Moy et al., 2018a; Jeevakumar et al., 2020; Shiers et al., 2020b) and our lack of knowledge concerning the actions of IL-6 on human DRG neurons. We developed an *in vitro* explant assay to study nascent protein synthesis in the DRG. Using this assay, we show that IL-6 causes nascent protein synthesis in human DRG neurons. This effect occurs in TRPV1+ neurons, demonstrating that it occurs across human nociceptors (Shiers et al., 2020a; Shiers et al., 2021; Tavares-Ferreira et al., 2022), and is blocked by eFT508, a specific inhibitor of MNK1/2 (Reich et al., 2018). We conclude that this signaling mechanism of IL-6 is conserved in human DRG neurons. Our work supports the targeting of translation regulation mechanisms for pain treatment in humans.

## METHODS

### Acute Human DRG Explants

Human tissue recovery was performed in accordance with pre-approved guidelines set by the Institutional Review Boards at the University of Texas at Dallas (UTD). In collaboration with the Southwestern Transplant Alliance (STA), lumbar human DRGs were recovered from neurologic determination of dead organ donors as described previously (Shiers et al., 2021; Tavares-Ferreira et al., 2022). Donor information is provided in **Table 1**. Human DRGs were transported to UTD in ice-cold, oxygenated artificial cerebrospinal fluid containing *N*-methyl-D-glucamine (NMDG-aCSF) as described previously (Valtcheva et al., 2016). After arrival, whole DRGs were immediately moved to ice cold, actively oxygenating NMDG-aCSF, cleaned, and then rapidly sliced into 1 mm explants. Slices were then distributed into treatment groups and transferred to actively oxygenating recording/labeling-aCSF (pH 7.4; 32.5 LJC) (R/L-aCSF) for 3.5 hrs or 4 hrs pre-treatment recovery.

**Table 1.** Organ Donor Demographics.

| DONOR | Tissue | Age | Sex | Race | COD | Figure | Datasets |
| --- | --- | --- | --- | --- | --- | --- | --- |
| 1 | L5 DRG | 37 | M | Black | CVA/Stroke | 1 | 1-4 |
| 2 | L5 DRG | 47 | F | White | Anoxia/Cardiovascular | 2 | 1-4 |
| 3 | L5 DRG | 28 | M | Hispanic | Anoxia/Cardiovascular | 3 | 1-4 |
| 4 | L4 DRG | 57 | F | White | Anoxia/Cardiac Arrest |  | 1-2 |
| 5 | L5 DRG | 19 | M | White | Asphyxiation/Suicide | 4 | 4 |
| 6 | L4 DRG | 20 | F | Hispanic | Head Trauma/MVA |  | 4 |

Formulas for aCSF solutions are provided in **Table 2**.

**Table 2.**
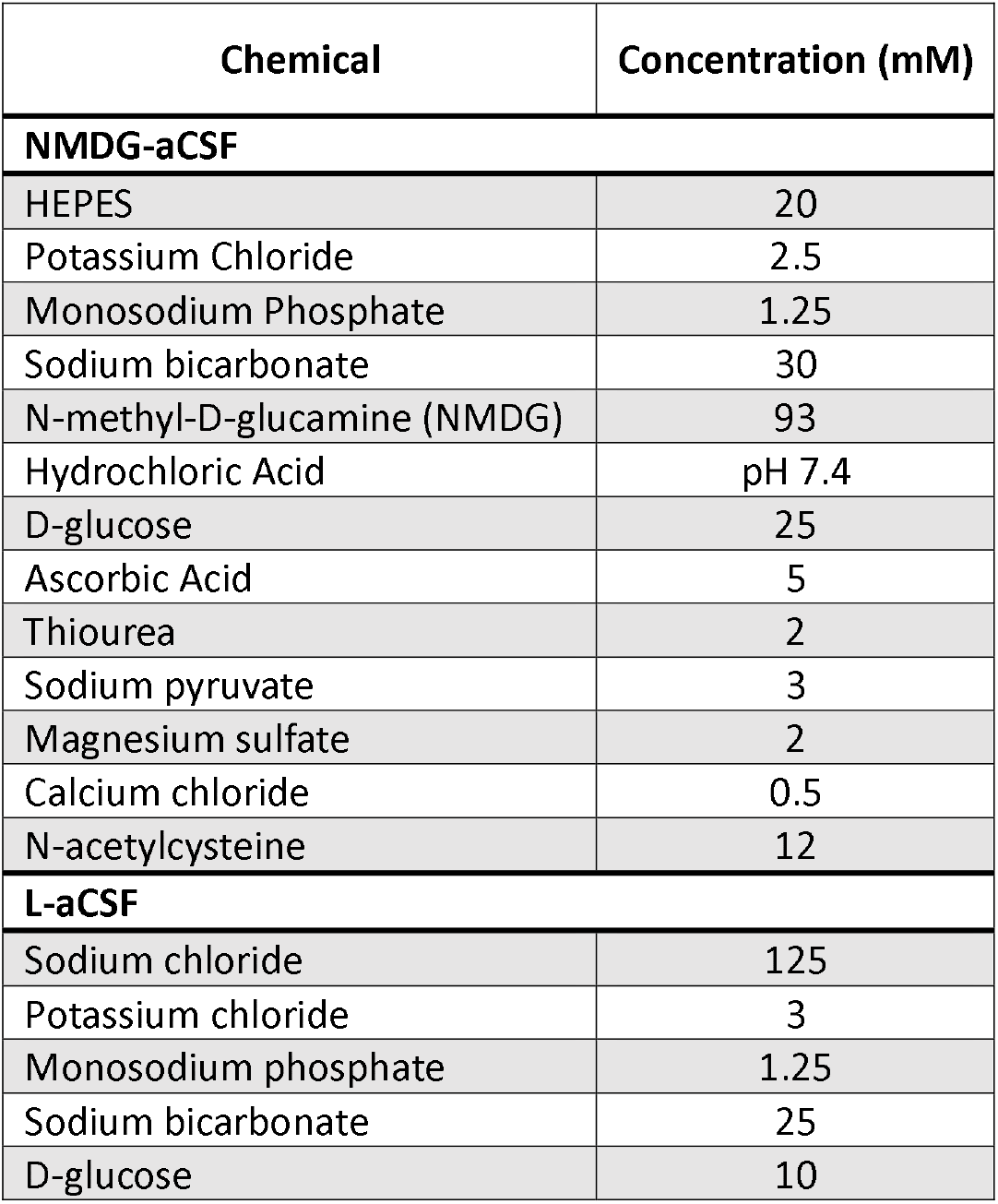
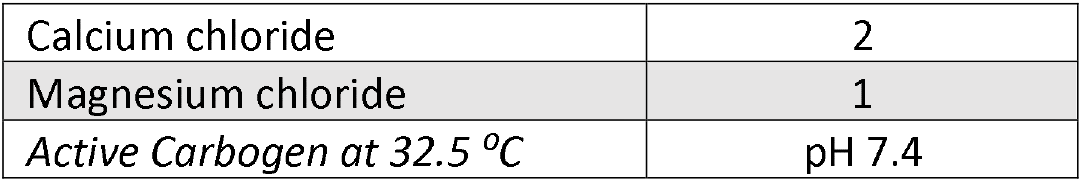
Artificial Cerebrospinal Fluid Formulas used in this study.

| Chemical | Concentration (mM) |
| --- | --- |
| **NMDG-aCSF** |  |
| HEPES | 20 |
| Potassium Chloride | 2.5 |
| Monosodium Phosphate | 1.25 |
| Sodium bicarbonate | 30 |
| N-methyl-D-glucamine (NMDG) | 93 |
| Hydrochloric Acid | pH 7.4 |
| D-glucose | 25 |
| Ascorbic Acid | 5 |
| Thiourea | 2 |
| Sodium pyruvate | 3 |
| Magnesium sulfate | 2 |
| Calcium chloride | 0.5 |
| N-acetylcysteine | 12 |
| **L-aCSF** |  |
| Sodium chloride | 125 |
| Potassium chloride | 3 |
| Monosodium phosphate | 1.25 |
| Sodium bicarbonate | 25 |
| D-glucose | 10 |
| Calcium chloride | 2 |
| Magnesium chloride | 1 |
| Active Carbogen at 32.5 °C | pH 7.4 |

### AHA Labeling and Drug Treatments

For all experiments in this study, L-Azidohomoalanine (AHA) (Click Chemistry Tools 1066) was used at 4 mM in R/L-aCSF (AHA-aCSF) for labeling of nascent proteins. Per Organ Donor replicate, the same base AHA-aCSF solution (vehicle, <0.5% DMSO) was used across treatment for 1 h incubations. For negative control, protein synthesis was inhibited using Anisomycin (60 µM; BioVision 1549). For protein synthesis stimulation, explants were treated with 10 ng/mL human recombinant IL- 6 and soluble IL-6 receptor (IL-6-sIL-6R) (1:1; R&D Systems 206-IL-050; I5771) for 20 mins. To inhibit IL-6-sIL-6R-induced MNK1/2 activity, explants were co-treated with 100 nM eFT508 during and after the pulse.

### FUNCAT and IF

Human DRG explants were fixed by rapid submersion into paraformaldehyde (PFA) (4%) and fixed for 15 min in pH 6.8 PFA and then to pH 9-10 PFA for an additional 20 min and then to pH 7.4 PFA for a final 25 min. Slices were then flash frozen in powdered dry ice and stored at -80 LJC until serial cryosectioning (80 µm: Donors 1-4; 100 µm: Donors 5-6) into free-floating sections for click chemistry. Incorporated AHA was detected using Fluorescent Non-canonical Amino Acid Tagging (FUNCAT) via cycloaddition of alkyne-conjugated Alexa Fluor 647 (Thermo Scientific A10278) (Dieterich et al., 2006; Dieterich et al., 2007). All click chemistry and other reagents used in this study are listed in **Table 3**.

**Table 3.**

| REAGENT or RESOURCE | SOURCE | IDENTIFIER |
| --- | --- | --- |
| **Antibodies** |  |  |
| Rabbit $\alpha$ TRPV1 | ThermoFisher | PA1-748 |
| Mouse IgG1 $\alpha$ NeuN | EMD Millipore | MAB377 |
| Rabbit $\alpha$ Phospho-S209-eIF4E | Abcam | Ab76256 |
| Goat $\alpha$ rabbit Alexa Fluor 488 | ThermoFisher | A11034 |
| Goat $\alpha$ mouse IgG1 Alexa Fluor 488 | ThermoFisher | A21121 |
| Goat $\alpha$ rabbit Alexa Fluor 555 | ThermoFisher | A21428 |
| DAPI |  |  |
| **Click Chemistry Reagents** |  |  |
| L-Azidohomoalanine (AHA) | Click Chemistry Tools | 1066 |
| Alkyne-Alexa Fluor 647, triethylammonium salt | ThermoFisher | A10278 |
| Tris(3-hydroxypropyl triazolyl-methyl)amine] (THPTA) | Click Chemistry Tools | 1010 |
| Sodium L-ascorbate | Sigma | A7631 |
| Copper(II) Sulfate | Sigma | C1297 |
| 1M Tris, pH 8.0 | ThermoFisher | AM9856 |
| Saponin | Sigma | 47036 |
| Bovine serum albumin (BSA) | ThermoFisher | SH3008803IR |
| Triton X-100 | Sigma | X100 |
| **Artificial cerebrospinal fluid (aCSF) for transport** |  |  |
| N-methyl-D-glucamine (NMDG) | Sigma |  |
| **Inhibitors and Interleukin-6-soluble receptor Pulse-Chase** |  |  |
| Dimethyl sulfoxide (DMSO) | Sigma | D8418 |
| Anisomycin | BioVision | 1549 |
| Human recombinant IL-6 | R&D Systems | 206-IL-050 |
| Human recombinant IL-6 receptor | R&D Systems | I5771 |
| eFT508 | MedChemExpress | HY-100022 |
| **Software and algorithms** |  |  |
| Olympus FLUOVIEW | Olympus | https://www.olympus-lifescience.com |
| FIJI (ImageJ) | NIH ImageJ | https://imagej.net/software/fiji/downloads |
| GraphPad Prism 10 | Dotmatics | https://www.graphpad.com/ |

Click chemistry reactions were carried out at room temperature while mixing for 20 to 24 hrs. Sections were then processed for immunofluorescence (IF) to detect either TRPV1 (488) or p-S209-eIF4E (488 or 555) and NeuN (488 or 555) (Shiers et al., 2021; Mitchell et al., 2023). All antibodies used in this study are listed in **Table 3**. Samples were treated with primary antibodies overnight at room temperature and incubated with secondary antibodies the next day for 1 hr before mounting.

### Fluorescence Intensity Imaging and Quantification of FUNCAT and p-eIF4E IF Signals

An Olympus FV3000RS laser scanning confocal microscope was used for imaging of human DRG sections. Four-color 10x images were acquired of entire human DRG sections for fluorescence intensity analyses. Two to 18 images were acquired for 7 to 20 sections per treatment condition for 6 Donor replicates. Acquisition parameters for FUNCAT and/or p-S209-eIF4E IF were kept consistent across experiments. Gain (1.000) and offset (4%) parameters were held constant for every section. Lipofuscin, a structure inherent to human nervous tissues that is highly autofluorescent, was detected as described previously for subtraction (Mitchell et al., 2023). For explant fluorescence intensity approximations, FUNCAT signal was measured across the field of view for the entire section surface area for 4 donors. For DRG neuronal somata, ROIs were identified in 12-bit images of TRPV1 IF (nociceptors) or NeuN IF (DRG neurons) channels. Somata were defined as having a minimum surface area of 350 µm^2^(Mitchell et al., 2023)1. Then, raw FUNCAT and/or p-eIF4E IF signals were measured per ROI, respectively.

### Subtraction and Transformation of Autofluorescent Lipofuscin Signal

Lipofuscin auto-signal was subtracted from each channel in 10x images using FIJI (NIH, Bethesda, MD). Auto-signal ROIs were identified using automated detection or manual tracing as described previously (Mitchell et al., 2023). Our automated macro parameters and codes are available upon request. Confocal images were then transformed by uniformly setting the lipofuscin signal (AU) to 1 AU. This permitted subtraction to mitigate its effect on FUNCAT and p-eIF4E IF intensity measurements per soma. This was done for all FUNCAT and p-eIF4E IF fluorescence intensities presented in this study using the following formula:

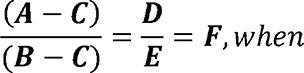

**A=** Integrated Density (AU·µm^2^), FUNCAT or p-eIF4E IF (raw)

**B=** Soma ROI Surface Area (µm^2^), TRPV1 IF or NeuN IF

**C=** Integrated Density (AU·µm^2^) of Lipofuscin (where AU= 1)

**D=** Measurable Integrated Density (AU·µm^2^) of FUNCAT or p-eIF4E IF

**E=** Measurable Signal Area (µm^2^) in soma ROI

**F=** Measurable FUNCAT or p-eIF4E IF signal (AU) in soma ROI

### Statistical Analyses

Graphs and Statistical analyses were generated using GraphPad Prism version 9.4 (GraphPad Software, Inc. San Diego, CA USA). All statistical tests used with associated biological replicate sizes, parameters, and *p* values are described in figure legends. Data in graphs is shown as dot plots for all data points and also represented as mean ± *SEM*. For both explant and somata analyses, transformed intensities were compared between treatment conditions either pairwise using *t-tests* where N= Donor Replicates, or for 3 or more conditions using Kruskal-Wallis with Dunn’s *post hoc* test with multiple comparisons where N= serial cryosections or neuronal somata. *P* values from multiple comparisons results are represented in all associated figures. All *p* values are expressed as \**p* <0.05; \*\**p* <0.005; \*\*\**p* <0.0005; \*\*\*\**p* <0.0001 in figures.

## RESULTS

### IL-6-sIL-6R Treatment Enhances Protein Synthesis in Human DRG Neurons by FUNCAT

We hypothesized that the IL-6-sIL-6R complex would induce nascent protein synthesis in human DRG tissue. To test this, we performed AHA labeling and FUNCAT in human DRG explants recovered from organ donors. We first sought to test whether FUNCAT fluorescence intensity reliably reports the presence of newly synthesized proteins. If this were true, then anisomycin treatment should reduce FUNCAT signal compared to vehicle baseline, serving as a negative control for the translation dependence of AHA incorporation (**Figure 1A-G**). We found anisomycin (60 µM) inhibited FUNCAT signal in cryosections from human DRG explants when compared to those treated with vehicle (**Figure 1B-C, E-F, H**). We next asked if treatment with the IL-6-sIL-6R complex enhanced FUNCAT signal when compared to vehicle baseline. We tested this by administering a 20-minute pulse of the complex (10 ng/mL) to explants. Our analysis revealed that IL-6-sIL-6R induces increased FUNCAT signal (**Figure 1D, G, and H**). These findings demonstrate that the IL-6-sIL-6R complex drives nascent protein synthesis in human DRG explants.

**Figure 1.**
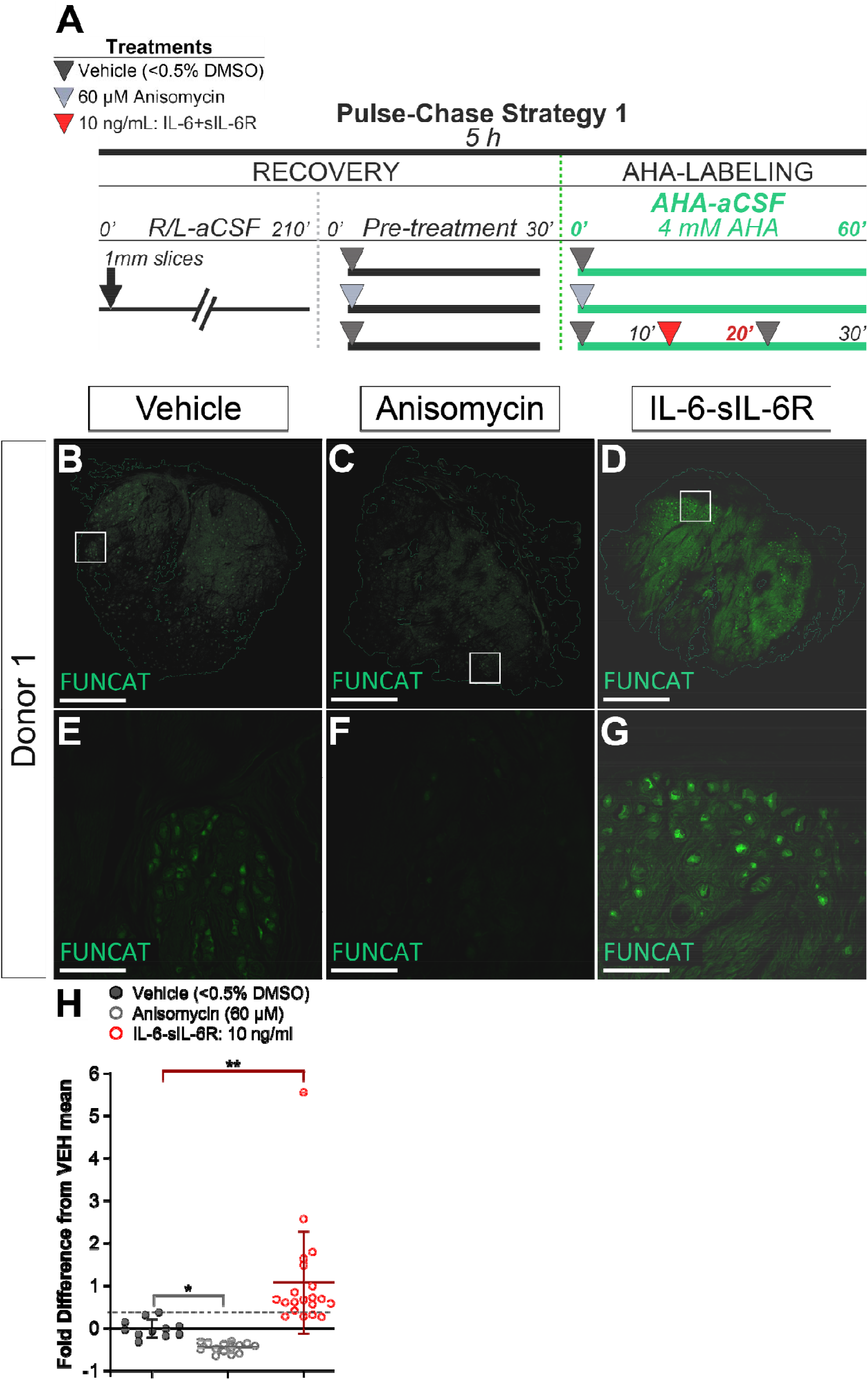
FUNCAT fluorescence intensity depends on protein synthesis and is increased after 20 min pulse with the IL-6-sIL6R complex in human DRG explants. (**A**) Pulse-Chase Treatment Strategy 1 for AHA-labeling of 1 mm human DRG explants for FUNCAT. Three parallel treatments per Donor: vehicle (<0.5% DMSO), baseline anisomycin (60 µM), or 20 min IL-6-sIL-6R pulse (10 ng/mL) for N=4 Donors. (**B-G**) Nascent proteins by FUNCAT in cryosections (80 µm) from human DRG explants treated with vehicle (**B, E**), anisomycin (**C, F**), or IL-6-sIL-6R (**D, G**) (Alkyne-Alexa-647, green). (**B-D**) 1.25x images of longitudinal DRG sections (scale bar, 2 mm). (**E-G**) Associated 10x images of DRG neuronal somata (scale bar, 500 µm). (**H**) Quantitation of FUNCAT intensity fold difference from vehicle baseline in cryosections from two technical replicates for four biological replicates. Scatter plot, values expressed as mean ± *SEM*. N= 80 µm serial sections across 1 mm explants from vehicle, N= 11; anisomycin, N= 14; IL-6-sIL-6R, N= 20. Violin plot, Kruskal-Wallis test with *post hoc* Dunn’s test: vehicle vs anisomycin, *p*= 0.0464; vehicle vs IL-6-sIL-6R, *p*= 0.0086. \**p* <0.05, ** *p* <0.01.

### IL-6-sIL-6R Treatment Induces Protein Synthesis in Human Nociceptors

We next set out to determine whether the IL-6-sIL-6R complex drives protein synthesis in human nociceptors. To test this, we treated human DRG explants with anisomycin, IL-6-sIL-6R, or vehicle as described above, but analyzed FUNCAT signal per nociceptor soma (**Figure 2**). To identify nociceptors in human DRG explants, we immunostained FUNCAT-treated sections for TRPV1 which is expressed by approximately 75% of human DRG neurons including all nociceptors (Shiers et al., 2020a; Shiers et al., 2021; Tavares-Ferreira et al., 2022). We then examined the colocalizing FUNCAT signal in TRPV1+ nociceptor cell bodies between conditions (**Figure 2A-C**). As expected, anisomycin dramatically inhibited baseline somatic FUNCAT signal in TRPV1+ nociceptors (**Figure 2A-E)**. Our analysis also revealed that IL-6-sIL-6R robustly increases FUNCAT signal within this subset of nociceptive neurons (**Figure 2A-E**). These findings show that the IL-6-sIL-6R drives nascent protein synthesis in human nociceptors.

**Figure 2.**
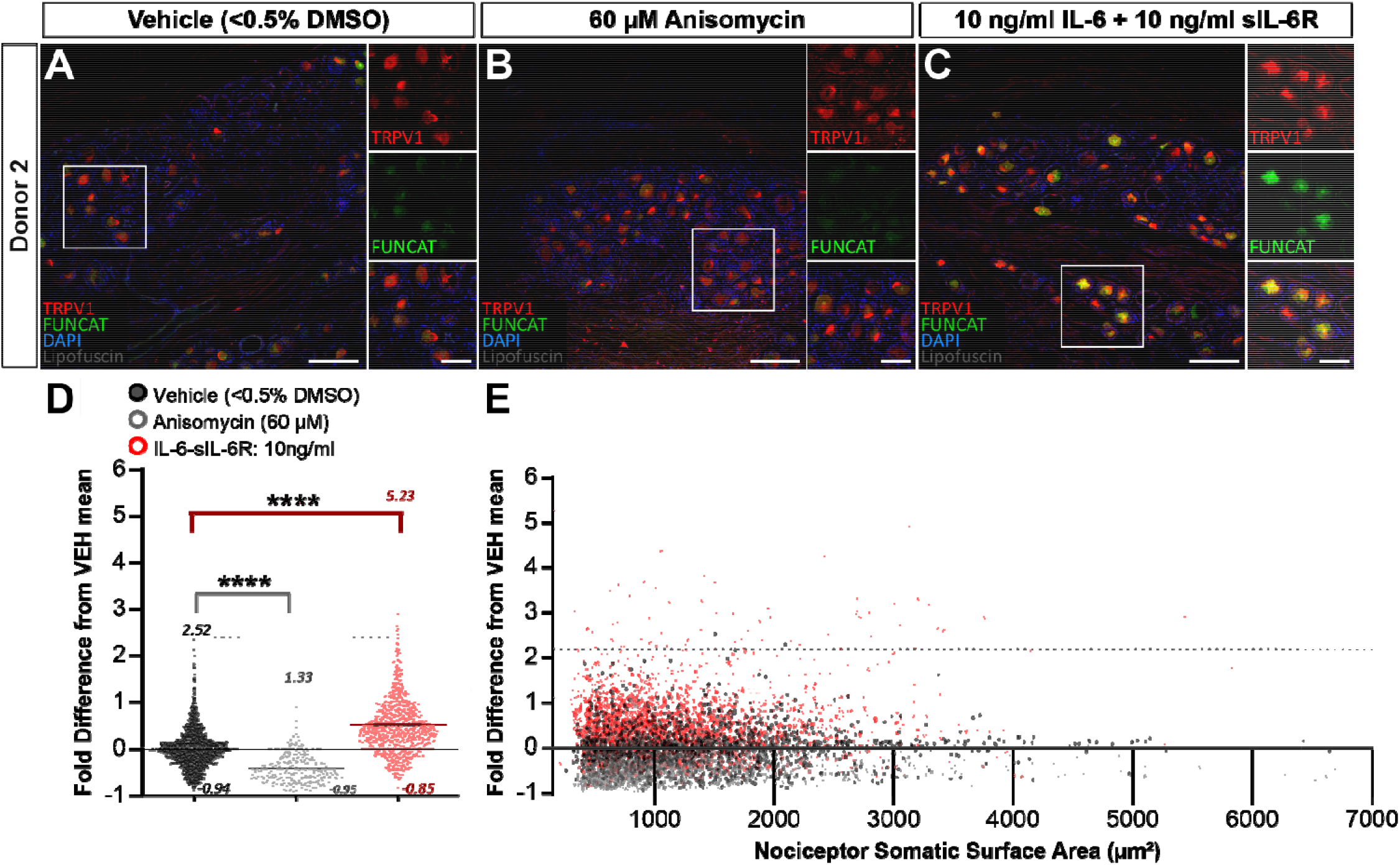
IL-6-sIL-6R stimulation enhances FUNCAT fluorescence intensity in a subset of human nociceptors. (**A-C**) Immunostaining for TRPV1 with FUNCAT and DAPI counterstain in cryosections (80 µm) from human DRG explants treated with vehicle (**A**), anisomycin (**B**), or IL-6-sIL-6R (**C**) (TRPV1, red; Alkyne-Alexa-647, green; DAPI, blue). 10x images (scale bar, 500 µm) with 5x magnified insets (scale bar, 100 µm). (**D**) Quantitation of somatic FUNCAT intensity fold difference from vehicle baseline in TRPV1+ nociceptors from two technical replicates for four biological replicates. Values expressed as mean ± *SEM*. \**p* <0.05; \*\*\*\**p* <0.0001. N= organ donor comparison of mean somata intensity difference. Total number of neurons: vehicle, N= 1,589; anisomycin, N= 1,357; IL-6-sIL-6R, N= 2,600. Kruskal-Wallis test with post hoc Dunn’s test: vehicle vs anisomycin, *p* <0.0001; vehicle vs IL-6-sIL-6R, *p* <0.0001. (**E**) Distribution of FUNCAT intensities detected in TRPV1+ nociceptors per somatic ROI surface area (µm^2^). Total TRPV1+ nociceptor somata comparison of fold intensity difference from vehicle baseline. N= TRPV1+ nociceptor somata from vehicle, vehicle, N= 1,589; anisomycin, N= 1,357; IL-6-sIL-6R, N= 2,600. XY Scatter plot, Kruskal-Wallis test with post hoc Dunn’s test: vehicle vs anisomycin, *p* <0.0001; vehicle vs IL-6-sIL-6R. *p* <0.0001.

### IL-6-sIL-6R Treatment Enhances p-eIF4E IF Signal in Human DRG Neurons

We next asked whether IL-6-sIL-6R treatment drives phosphorylation of eIF4E at serine residue 209 (p-eIF4E) (**Figure 3**). We tested whether IL-6-sIL-6R enhances this signaling by immunostaining for p-eIF4E in sections from human DRG explants treated as described above. We then analyzed p- eIF4E immunofluorescence (p-eIF4E IF) signal per DRG neuronal soma (**Figure 3 A-B**). We found that p-eIF4E IF was increased after IL-6-sIL-6R treatment when compared to vehicle baseline (**Figure 3C**). This change in signaling was also readily evident in our images where more than 50% of neurons appeared to exhibit dramatically enhanced p-eIF4E IF intensity (**Figure 3D**). The data demonstrate that phosphorylation of eIF4E at serine 209 is induced by IL-6-sIL-6R-signaling in human DRG neurons. This suggests that this signaling pathway may be responsible for increased nascent protein synthesis in IL-6-sIL-6R-treated hDRG neurons, a hypothesis we tested in a subsequent experiment.

**Figure 3.**
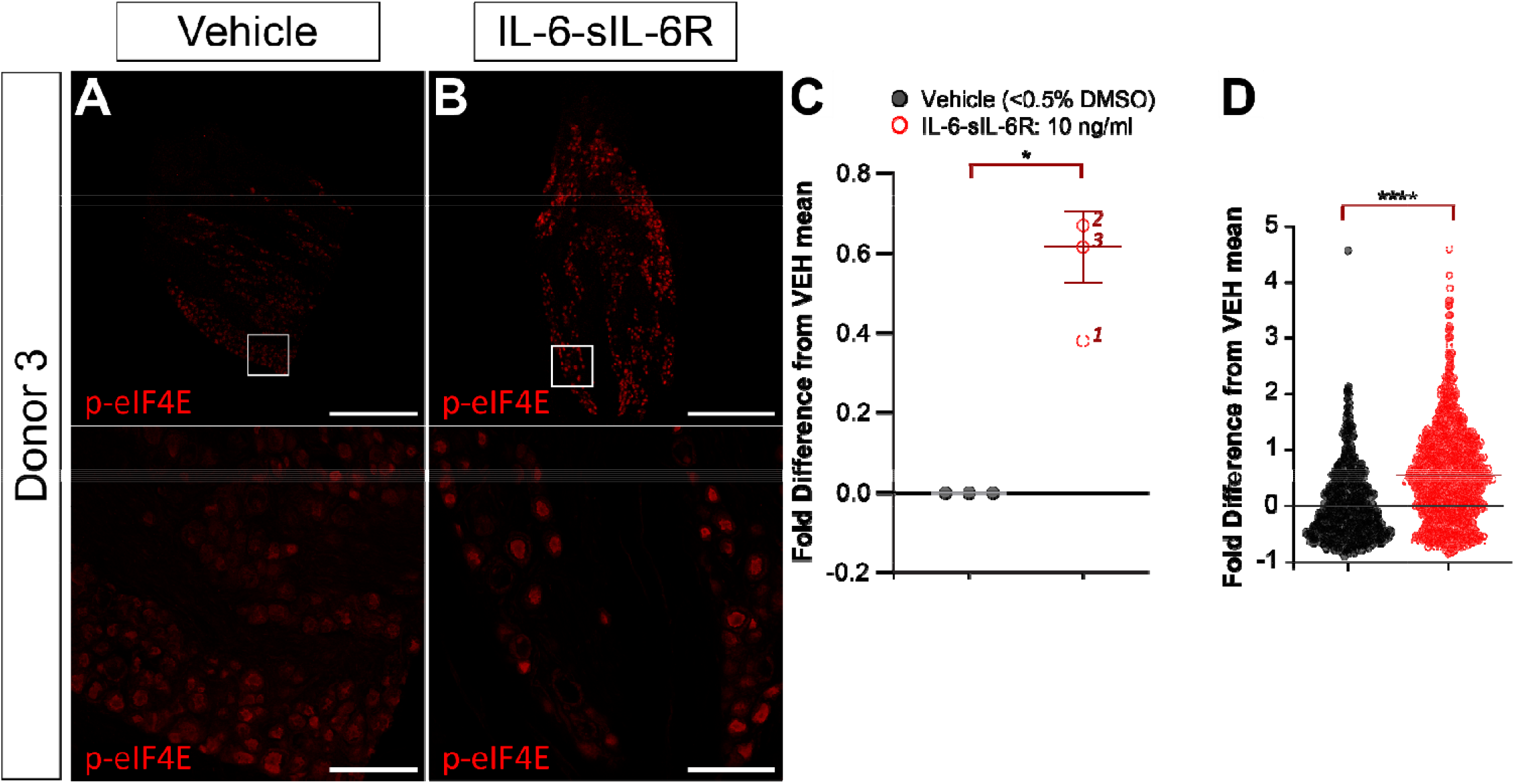
IL-6 stimulation enhances p-eIF4E immunofluorescence intensity in a subset of human DRG neurons. (**A-B**) Immunostaining for p-eIF4E (phosphor-S209) in cryosections (80 µm) from human DRG explants treated with vehicle (**A**) or IL-6-sIL-6R (**B**) (p-eIF4E, red).1.25x images of longitudinal DRG sections (scale bar, 2 mm). Insets show 10x images of human DRG neurons (scale bar, 500 µm). (**C**) Quantitation of somatic p-eIF4E IF intensity fold difference from vehicle baseline in DRG neurons in cryosections from two technical replicates for three biological replicates. N= 3 organ donor comparison of mean intensity difference after IL- 6-sIL-6R-treatment. Scatter plot, paired *t*-test, two-tailed, *p*= 0.0249. (**D**) Spread of fold IF intensity difference from vehicle baseline per DRG neuron. N= DRG neuronal somata from vehicle, N= 677; IL-6-sIL-6R, N= 921. Violin plot. Kruskal-Wallis test with post hoc Dunn’s test, *p* <0.0001.

### IL-6-sIL-6R Treatment Drives MNK1/2-Dependent Phosphorylation of eIF4E and Protein Synthesis in Human DRG Neurons

We sought to determine whether MNK1/2 activity is required for IL-6-sIL-6R-driven increases in p- eIF4E IF and FUNCAT signals in human DRG neurons (**Figure 4**). To test this possibility, we u ed the specific MNK1/2 inhibitor eFT508 (100 nM) as a co-treatment with 20 min IL-6-sIL-6R pulse (**Figure 4A**). If IL-6-sIL-6R-driven enhancements in p-eIF4E and nascent proteins in human DRG neur ns depend on MNK1/2, then eFT508 should block increases in p-eIF4E IF and FUNCAT in these neurons. IL-6-sIL-6R treatment increased p-eIF4E IF and FUNCAT signals in human DRG neurons when compared to vehicle (**Figure 4B-F**). Co-treatment with eFT508 inhibited IL-6-sIL-6R-driv n increases in p-eIF4E IF signal below normal baseline in human DRG neurons (**Figure 4E**). Co-treatment with eFT508 also reduced IL-6-sIL-6R treatment-induced increases in FUNCAT to baseline intensities (**Figure 4F**). We sought to investigate variation in p-eIF4E IF and FUNCAT signals across individual neuronal soma compared to vehicle treatment. As shown in **Figures 4E and 4F**, plotting our intensity data as a function of soma surface area size revealed that many individual neurons w thin the IL6-sIL-6R-treated explants exhibited increased p-eIF4E signals when compared to vehicle, while a large portion of those in the eFT508-cotreated explants displayed intensities below baseline (**Figure 4G**). Similarly, analysis of FUNCAT signal in IL6-sIL-6R-treated explants showed notable increases compared to vehicle, whereas eFT508-cotreated explants did not exhibit similar decreases (**Figure 4H**). Our quantitative data appear to indicate clustering towards smaller neuronal sizes for both p- eIF4E IF and FUNCAT; however, by using soma surface area, we did not account for well-known effects of intracellular fluorescent dye concentration on intensity detection which is a potential confound of the result (Fink et al., 1998; Waters, 2009; Morikawa et al., 2016). Again, FUNCAT signal was not decreased below baseline with eFT508 treatment. Taken together, the data demonstrates that IL6-sIL-6R treatment increased nascent protein synthesis in human DRG neurons via MNK- eIF4E signaling. Critically, engagement of this pathway is directly correlated with nascent protein synthesis on a per-neuron basis in human DRG.

**Figure 4.**
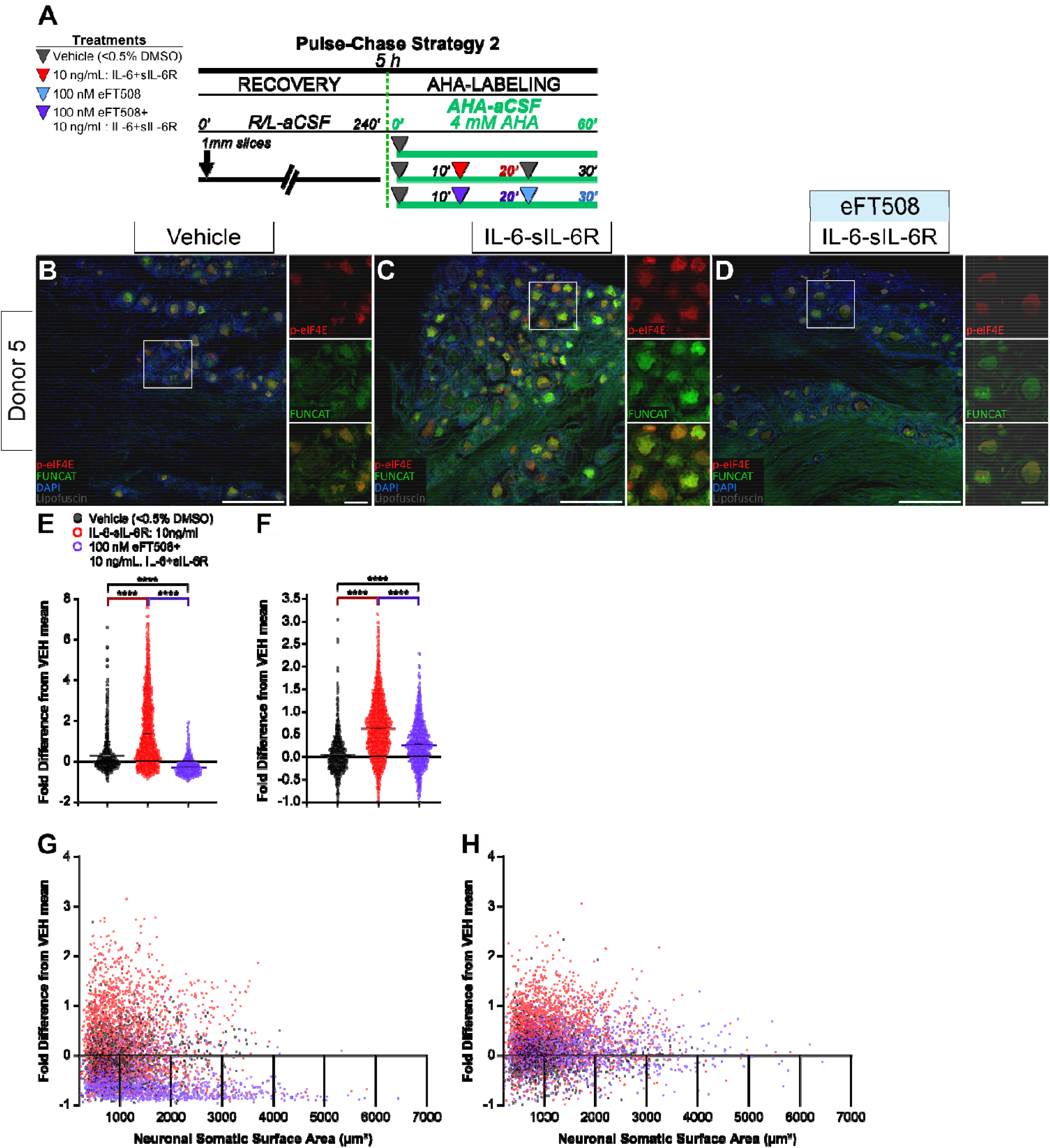
IL-6-stimulated increases in p-eIF4E IF and FUNCAT fluorescence intensity depends on MNK1/2 activity. (**A**) Pulse-Chase Treatment Strategy 2 for AHA-labeling of 1 mm human DRG explants. Three parallel treatments per Donor: vehicle (<0.5% DMSO), 20 min IL-6-sIL-6R pulse, or 100nM eFT508 with 20 min IL-6-sIL-6R pulse. (**B-D**) Immunostaining for p-eIF4E with FUNCAT and DAPI in cryosections (100 µm) from human DRG explants treated with vehicle (**B**), IL-6-sIL-6R (**C**), or eFT508 with IL-6-sIL-6R (**D**) (p-eIF4E, red; Alkyne-Alexa-647, green; DAPI, blue). 10x images (scale bar, 500 µm) with 5x magnified insets (scale bar, 100 µm). (**E-F**) Quantitation of somatic p-eIF4E IF (**E**) or FUNCAT (**F**) intensity fold difference from vehicle baseline in DRG neurons. Three technical replicates for 5 Donors (vehicle and IL-6-sIL-6R) and 2 Donors for (eFT508+ IL-6-sIL-6R), N= DRG neurons from vehicle, N= 2,263; IL-6-sIL-6R, N= 5,250; eFT508+IL-6-sIL-6R, N=2,395. Violin plot, values expressed as mean ± *SEM*. \*\*\*\**p* <0.0001. Kruskal-Wallis test with post hoc Dunn’s test. (**G-H**) Distribution of p-eIF4E IF intensity (**G**) or FUNCAT intensity (**H**) detected in neurons per somatic ROI surface area (µm^2^). Total neuronal somata comparison of fold intensity difference from vehicle baseline. Three technical replicates for 5 Donors (vehicle and IL-6-sIL-6R) and 2 Donors for (eFT508+ IL-6-sIL-6R), N= DRG neurons from vehicle, N= 2,263; IL-6- sIL-6R, N= 5,250; eFT508+IL-6-sIL-6R, N=2,395.

## DISCUSSION

Activity-dependent translation of new proteins has long been recognized as a core mechanism for synaptic plasticity in the CNS (Kelleher et al., 2004; Costa-Mattioli et al., 2009) and for changes in nociceptor excitability in the PNS (Khoutorsky and Price, 2018; Yousuf et al., 2021). Our foundation of knowledge on this key mechanism for neuronal plasticity is almost entirely built on experiments done in rodents, largely mice. In this body of work, we demonstrate that IL6-signaling causes an activity-dependent increase in nascent protein synthesis in human DRG neurons recovered from organ donors. This effect is entirely dependent on MNK signaling because it is blocked by eFT508, a specific inhibitor of both MNK1 and MNK2 (Reich et al., 2018), both of which are expressed by human DRG neurons (Shiers et al., 2023). The increase in nascent protein synthesis is also associated with increased p-eIF4E, a specific target of MNK1/2 signaling (Pyronnet et al., 1999), providing biochemical evidence of engagement of this pathway in human DRG neurons. Finally, our findings show that IL-6-sIL-6R-induced increases in nascent protein synthesis occur in TRPV1+ neurons in human DRG, demonstrating that this signaling pathway is engaged in human nociceptors (Shiers et al., 2020a; Shiers et al., 2021; Tavares-Ferreira et al., 2022). We conclude that this signaling mechanism, which is directly associated with IL-6-induced nociception in mice (Melemedjian et al., 2010; Moy et al., 2017), is similarly engaged in humans supporting the conclusion that IL-6-induced pain in humans is also driven by MNK activation in nociceptors.

An important observation in our experiments is that MNK signaling only appears to be required for IL- 6-sIL-6R-enhanced protein synthesis and not for constitutive protein synthesis in human DRG neurons. Treatment with eFT508 clearly decreased p-eIF4E levels below baseline in our experiments but it did not suppress baseline AHA-incorporation. In contrast, anisomycin, which blocks peptide bond formation, decreased AHA-incorporation well below baseline levels. The effect of eFT508 treatment in our experiments was specific for blocking the enhanced AHA-incorporation, measured as FUNCAT signal, when slices were co-treated with IL-6-sIL-6R. It has long been recognized that MNK-eIF4E signaling influences the translation of a subset of mRNAs that are involved in neuronal plasticity (Aguilar-Valles et al., 2018; Amorim et al., 2018; Moy et al., 2018b; Megat et al., 2019), immune response (Joshi et al., 2009), and oncogenesis (Furic et al., 2010; Altman et al., 2013). Some of these mRNAs have been identified in cancer cells, immune cells and in DRG neurons (Furic et al., 2010; Aguilar-Valles et al., 2018; Amorim et al., 2018; Moy et al., 2018b), but the guiding principles for how p-eIF4E regulates the translation of specific subsets of mRNAs have still not been discovered (Scheper and Proud, 2002; Chen et al., 2023). Given the importance of this pathway for pain, and our demonstration that it is engaged in human nociceptors, a future priority should be using techniques like ribosome profiling to understand precisely which mRNAs are translated when p-eIF4E levels are increased in human nociceptors.

We recently demonstrated that blocking MNK signaling in human DRG neurons recovered from patients undergoing thoracic vertebrectomy surgery reverses spontaneous action potential activity that is associated with neuropathic pain symptoms in these patients (Li et al., 2023). Previous RNA sequencing experiments on DRGs from these patients found that cytokines and chemokines are upregulated in DRGs from male and female patients who had neuropathic pain in associated dermatomes (North et al., 2019; Ray et al., 2023). Our work with IL-6-sIL-6R links these electrophysiological findings in patient DRG neurons to induction of this signaling pathway using a system that can be used to screen other cytokines or chemokines or even groups of cytokines and chemokines to examine whether MNK-eIF4E signaling is engaged by these factors. It can also be used in conjunction with proteomic or transcriptomic methods to identify the nascent proteins that are synthesized or the mRNAs that are translated, respectively. We envision that the assay described here can be used as a discovery platform for gaining a deeper understanding of the intricacies of human nociceptor plasticity.

## Author Contributions

TJP designed the research project; MEM, GT LFC, JMM, and LH conducted experiments; MEM, GT LFC, and SS performed imaging analysis; MEM and SS established imaging methods; MEM, GT and TJP wrote the paper. All authors approved the final version of the paper.

## Competing Interest Statement

TJP is a founder of 4E Therapeutics, a company developing MNK inhibitors for pain treatment. The authors declare no other financial conflicts of interest related to this work.

### Acknowledgments

The authors thank Erin Vines, Anna Cervantes, Geoffrey Funk, and Peter Horton at the Southwest Transplant Alliance. The authors are grateful to the organ donors and their families for their enduring gift that made this work possible. This work was supported by NIH grant U19NS130608 and R01NS111929 to TJP.

